# Tonotopic organization of auditory cortex in awake marmosets revealed by multi-modal wide-field optical imaging

**DOI:** 10.1101/2024.04.05.588237

**Authors:** Xindong Song, Yueqi Guo, Chenggang Chen, Jong Hoon Lee, Xiaoqin Wang

## Abstract

Tonotopic organization of the auditory cortex has been extensively studied in many mammalian species using various methodologies and physiological preparations. Tonotopy mapping in primates, however, is more limited due to constraints such as cortical folding, use of anesthetized subjects, and mapping methodology. Here we applied a combination of through-skull and through-window intrinsic optical signal imaging, wide-field calcium imaging, and neural probe recording techniques in awake marmosets (*Callithrix jacchus*), a New World monkey with most of its auditory cortex located on a flat brain surface. Coarse tonotopic gradients, including a recently described rostral-temporal (RT) to parabelt gradient, were revealed by the through-skull imaging of intrinsic optical signals and were subsequently validated by single-unit recording. Furthermore, these tonotopic gradients were observed with more details through chronically implanted cranial windows with additional verifications on the experimental design. Moreover, the tonotopy mapped by the intrinsic-signal imaging methods was verified by wide-field calcium imaging in an AAV-GCaMP labeled subject. After these validations and with the further effort to expand the field of view more anteroventrally in both windowed and through-skull subjects, an additional putative tonotopic gradient was observed more rostrally to the area RT, which has not been previously described by the standard model of tonotopic organization of the primate auditory cortex. Together, these results provide the most comprehensive data of tonotopy mapping in awake primate species with unprecedented coverage and details in the rostral proportion and supports a caudorostrally arranged mesoscale organization of at least three repeats of functional gradients in the primate auditory cortex, similar to the ventral stream of primate visual cortex.

## INTRODUCTION

The tonotopic organization is considered a fundamental feature of the auditory cortex in mammals. It has been widely demonstrated in the primary auditory cortex and reported also in many higher-order auditory cortical areas. Recent advances in recording techniques, particularly optical imaging, have facilitated a more comprehensive understanding of these features in small mammals (Issa, et al., 2014) (Romero, et al., 2020) (Narayanan, et al., 2023) (Kalatsky, et al., 2005) (Nelken, et al., 2004) (Langner, et al., 2009). In both rodents and carnivores, the primary auditory field A1 was found tonotopically organized, with its high-frequency end pointing anteriorly and adjoining other auditory fields (e.g., AAF) (Kaas, 2011) (Stiebler, et al., 1997) (Guo, et al., 2012) (Issa, et al., 2014) (Romero, et al., 2020) (Narayanan, et al., 2023) (Kalatsky, et al., 2005) (Nelken, et al., 2004) (Langner, et al., 2009). Conversely, in primates, early investigations using acute electrophysiological recordings identified the primary auditory field A1 shared a low-frequency tonotopic reversal with its more anterior neighbor (rostral field or R) (Merzenich & Brugge, 1973) (Imig, et al., 1977). These findings imply an early departure in evolution of primates from other mammals in terms of how the auditory cortex is functionally organized. Later studies expanded mapping coverage and suggested the existence of another tonotopically organized field located more anteriorly to R. This field was named the rostrotemporal field (RT) and is also considered as a part of the auditory core, exhibiting primary-like features (Morel & Kaas, 1992). The tonotopic organization of RT has not been studied adequately, comparing to those of A1 and R. Nevertheless, studies suggest that RT’s caudal end, contiguous with R, represents high frequencies (Kaas & Hackett, 2000) (Petkov, et al., 2006) (Bendor & Wang, 2008) (Joly, et al., 2014) (Baumann, et al., 2015) (Nishimura, et al., 2018) (Tani, et al., 2018) (Song, et al., 2022). Surrounding the auditory core, the secondary auditory cortex, also called auditory belt, contains several sub-fields in each of its lateral, medial and caudal parts. Both the lateral and the medial auditory belts, were found to have tonotopic gradients that are in parallel with their immediately neighboring field in the auditory core. For example, the mediolateral belt (ML) parallels A1, anterolateral belt (AL) parallels R, and possibly rostrotemporal lateral belt (RTL) parallels RT as well (Merzenich & Brugge, 1973) (Imig, et al., 1977) (Morel & Kaas, 1992) (Rauschecker, et al., 1995) (Kaas & Hackett, 2000) (Petkov, et al., 2006) (Kusmierek & Rauschecker, 2009). The caudal belt may also contain sub-fields that are tonotopically organized, although the evidence is generally much weaker and somewhat illusive than other fields (Kaas & Hackett, 2000) (Petkov, et al., 2006). In contrast to auditory core, where a majority of neurons are responsive to pure tones but not to broadband sounds, the auditory belt contains neurons that generally prefer broadband sounds (Rauschecker, et al., 1995) (Rauschecker & Tian, 2004) or harmonic sounds (Kikuchi, et al., 2014) over pure tones, implying a spectral processing hierarchy within the auditory pathway. Lateral to the auditory belt is the auditory parabelt, which is considered the tertiary auditory cortex due to its minimal direct connectivity with the auditory core but connectivity with the belt. The amount of physiological data in this area is very limited comparing to that in the core and belt. Only until recently, electrophysiological recordings in awake macaques and marmosets have uncovered significant number of neurons in parabelt that are sensitive to pure tones (Kajikawa, et al., 2015) (Gamble, 2020). The macaque data may further support a possible existence of tonotopic gradients in the parabelt, whereas marmoset data showed a clear high-frequency sensitive region in the caudal end of the rostral parabelt (RPB). However, the limited area coverage of these awake electrophysiological recordings hinders direct comparison across areas and cortical stages. On the other hand, optical imaging in anesthetized primates provide a broader area coverage and some clues about the tonotopic organizations in the higher-order regions (Nishimura, et al., 2018) (Tani, et al., 2018). But the anesthesia used in these studies makes it harder to interpret results in the higher order regions. Recently, we developed a through-skull intrinsic signal imaging technique that significantly broadens recording coverage in awake marmosets (Song, et al., 2022). Utilizing a phase-encoded mapping paradigm, a continuous low-frequency to high-frequency gradient from RT to RPB was discovered beyond conventionally described tonotopy in all tested hemispheres, indicating the need for further revision to the current model of primate auditory cortex (Kaas & Hackett, 2000) (Kaas, 2011). Furthermore, orthogonal to the well-established auditory hierarchy along the mediolateral direction, a recent anatomical study in macaques has demonstrated another caudorostrally directed pathway comprising step-wise projections from AI through R and RT, continuing to a more rostrally located rostrotemporal polar field (RTp) (Scott, et al., 2017). This pathway supports a rostrally directed flow of auditory processing similar to the ventral stream of macaque visual cortex. Nevertheless, the functional organization including tonotopic features at the rostral side of this pathway is largely unknown. Here, we seek to utilize optical imaging techniques to draw a more complete picture of tonotopic organizations in marmoset cortex.

One limitation of our previous through-skull imaging study and some other relevant studies is the lack of systematic cross-validation with other experiment design paradigms and functional recording modalities. To address this, we first validated our previous through-skull imaging results with subsequently performed single-unit recordings in three awake subjects. The imaged frequency tunings were found to be significantly correlated with the pooled single-unit frequency tunings, suggesting the imaging results can be explained by the underlying neuronal activities. In our previous through-skull imaging study we provided evidence that the tonotopic maps were very similar when imaged through-skull or through a subsequently implanted cranial window by the same phase-encoded experimental paradigm (Song, et al., 2022). The phase-encoded paradigm has been demonstrated to be a powerful tool to map continuous topographical gradient and can significantly reduce the experimental time required to acquire functional maps (Kalatsky & Stryker, 2003) (Nelken, et al., 2004) (Kalatsky, et al., 2005) (Joly, et al., 2014) (Baumann, et al., 2015). To compare it with a standard trial-based paradigm, we have performed both in a window-implanted subject. The results showed the phase-encoded paradigm can acquire a similar map as inferred by the trial-based paradigm, but with less time spent and more continuous coverage on tested feature dimension. Furthermore, the wide-field imaging we have performed so far was based on the intrinsic optical signal which is believed to reflect hemodynamic changes. Although it does not require any extrinsic labeling, the signal is slow and only measures indirect neural activities. To measure more direct neuronal activities using wide-field optical imaging, in one of our window-implanted subjects, we have performed adeno-associated virus (AAV) labeling for the calcium signal (AAV1-CaMKII-cre and AAV(DJ)-FLEX-GCaMP6s) (Song, et al., 2022). This virus combination presumably labels excitatory neurons’ activities. Further wide-field calcium fluorescence imaging showed that the signal is much faster and more sensitive than the intrinsic optical signal. A calcium trial-based tonotopic map was acquired and compared to the intrinsic phase-encoded map. These two maps were found to be highly correlated, suggesting the validity of mapping tonotopy with both signals. The wide-field calcium imaging demonstrated here may further enhance our ability to map the primate cortex and accelerate our understanding on how primate cortex is organized mesoscopically.

After all these validations, we performed intrinsic optical signal imaging in additional subjects with further effort to extend the FOV coverage more rostrally in both windowed and through-skull imaged subjects, since our preliminary results implied the possibility of additional tonotopically organized regions existing along this anatomical direction beyond the classical auditory regions. Our results revealed the presence of an additional tonotopic gradient more rostral to the RT core field, which runs roughly perpendicularly to the lateral sulcus (LS). The low-frequency end of this gradient is located closer to the LS and high-frequency end is located further from the LS. The location of this additional gradient presumably overlaps with the recently defined region rostro-temporal polar (RTp) (Scott, et al., 2017). This gradient has weaker tonal response and less salient tuning than more caudal gradients, a phenomenon similar to the belt versus core comparison. The existence of such a functional gradient, together with previous anatomical and connectivity evidence, supports the idea that the primate auditory cortex has a caudo-rostrally arranged information flow for hierarchically processing what in the sound that parallels ventral pathway of the primate visual cortex. In summary, the results demonstrate that the primate auditory cortex may be more complex than previously thought.

## RESULTS

### Verifying through-skull imaged tonotopic maps with electrophysiological recordings

To validate our through-skull imaging, we performed electrophysiological recording in three marmoset subjects after we obtained through-skull imaging data from the same subjects (Figure 1). In all three subjects (Figure 1A), the tonotopy can be clearly mapped by through-skull imaging with a phase-encoded experiment paradigm and Fourier-based analysis (Song, et al., 2022) – the posterior (and caudal-most) low frequency region marks the border between R and A1, while the more anterior (and also more rostral) low frequency region marks the rostral end of RT. Consistent with our previous observation, an unconventional low-to-high tonotopic gradient extending laterally and caudally from the low-frequency end of RT towards a high-frequency region outside the core was evident in all three subjects. The high-frequency region of this gradient has been shown to reside beyond the lateral belt and into the parabelt, presumably RPB, according to the registration to two different marmoset atlases (Song, et al., 2022) (Liu, et al., 2018) (Majka, et al., 2021).

**Figure 1.**
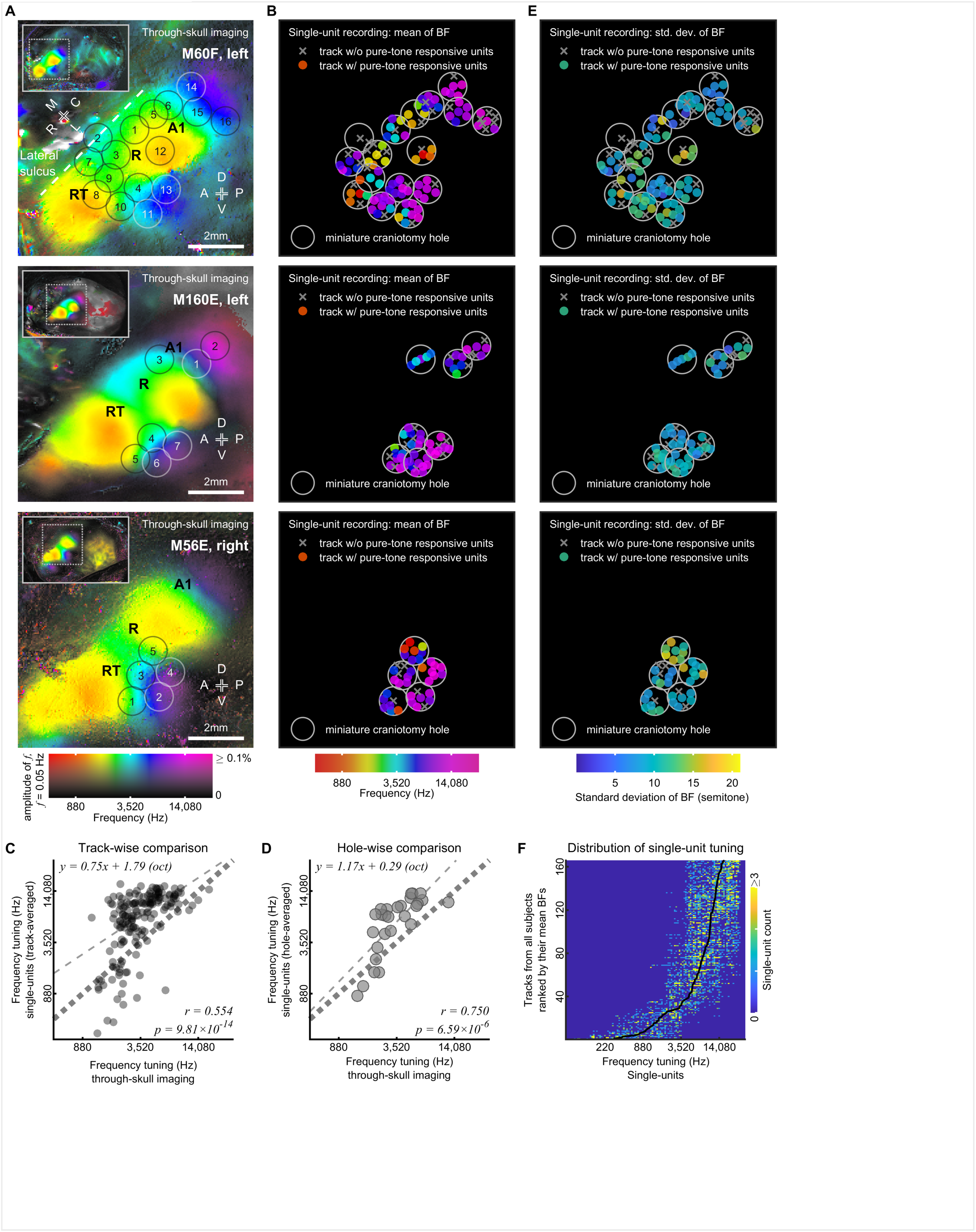
Verifying through-skull imaged tonotopy with electrophysiology. A. Tonotopic map obtained by through-skull intrinsic imaging in each tested subject (M60F, M160E, and M56E, from top to bottom). The locations of the three sub-fields of the auditory core are indicated by their names (A1, R, and RT). The locations of the subsequent craniotomy holes for electrophysiological recordings are labeled by black or white circles and numbered in their recording sequence. Inset: a zoomed-out view that covers the entire imaging chamber. Colormap: the 2D color code for visualizing the tone-tuning frequency (color) and the response amplitude (intensity) together for each pixel. Scalebar: 2mm. A: anterior; P: posterior; D: dorsal; V: ventral. R: rostral; C: caudal; M: medial; L: lateral. B. Mean best frequency of each recording track in the 64-channel single-unit recording. Colored dots: recording tracks with the color representing the mean best frequency (averaged on a logarithmic scale) of all units recorded in that track. Gray crosses: recording tracks that have no pure-tone responsive units. Grey circles: 1-mm miniature craniotomy holes for electrophysiological recording. C. Track-wise comparison of the frequency tunings between through-skull imaging and electrophysiology (n = 164). Dashed line: a linear fit of the data, with a slope of 0.75. The correlation coefficient (r) between the two measures is 0.554. D. Hole-wise comparison of the frequency tunings between through-skull imaging and electrophysiology (n = 27). Dashed line: a linear fit of the data, with a slope of 1.17. The correlation coefficient (r) between the two measures is 0.750. E. Variability of single-unit best frequencies in each recording track in the 64-channel single-unit recording. Colored dots: recording tracks with the color representing the standard deviation of the best frequencies of all pure-tone responsive units recorded in that track. Gray crosses: recording tracks that have no pure-tone responsive units. Gray circles: 1-mm miniature craniotomy holes for electrophysiological recording. Colormap: standard deviation on a logarithm scale (semitone). F. Distribution of single-unit tuning for each track. Each row contains a histogram of the single-unit tuning of a track, and the 164 rows are ranked by the mean BF of the track. The color indicates the count of the units within the BF range defined by a bin on the x-axis, with a saturation value of 3 counts. Black line: the mean BF of each track.

After through-skull imaging, subsequent multi-channel single-unit recording was performed through miniature craniotomy holes (Figure 1A and B). Each colored dot in Figure 1B represents a recording track, with the color representing the mean best frequency (BF) of recorded units in that track. The electrophysiology data in these subjects were mainly collected for another experiment and thus was biased towards covering mid-to-high frequency regions, especially for subjects M160E and M56E. Nevertheless, many features in the through-skull maps were confirmed by electrophysiology. Specifically, in subject M60F, the frequency tunings across the tracks in holes #12, #5, #6, #15, #14 and #16 show a tonotopic gradient that is in line with the tonotopic gradient in A1 estimated by the through-skull imaging data, while the tunings in holes #12, #3, and #7 are consistent with the estimated tonotopic gradient in R. Additionally, the tunings across the tracks in holes #8, #4, #10, #11 and #13 are also in general agreement with the recently described tonotopic gradient extending from the low-frequency RT to the putative parabelt (Song, et al., 2022). Furthermore, the non-responsive tracks in holes #2 and #7 are in good consistency with the location of the estimated lateral sulcus. In the second subject M160E, the tunings in holes #3, #1, and #2 show a mid-to-high tonotopic gradient that is in line with the tonotopic gradient in A1 estimated by through-skull imaging. Moreover, the tunings in holes #4 to #7 form another mid-to-high tonotopic gradient that is in general agreement with the recently described RT-RPB tonotopic gradient mapped by through-skull imaging. In the third subject M56E, the tunings in holes #1 to #4 are consistent with the high-frequency end of the RT-RPB tonotopic gradient mapped by through-skull imaging. The tunings in hole #5 show more variance, as the through-skull map indicates more dramatic tuning transitions around this location.

To further quantify the tuning relationship between the two methods, the frequency tunings derived from through-skull intrinsic signal imaging were compared with the track-averaged tuning (Figure 1C) and hole-averaged tuning (Figure 1D) from electrophysiology. The tunings between imaging and electrophysiology are significantly correlated (r = 0.554 for track-wise correlation, with p=-9.81×10^-14^; r = 0.750 for hole-wise correlation, with p=6.59×10^-6^). Linear models were fit to the data, with slope values of 0.75 (track-wise, Figure 1C) and 1.17 (hole-wise, Figure 1D). The total variance explained by the linear model increased from 30.7% in the track-wise comparison to 56.3% in the hole-wise comparison. This is consistent with the fact that the distance of the tracks from the mean center within the same hole is averaged as 0.28 mm, very close to the marmoset through-skull imaging resolution (0.30-0.35 mm, half-width at half-maximum [HWHM]) (Song, et al., 2022). Averaging single-unit tunings across each hole would thus better match the spatial diffusion scale of marmoset through-skull imaging. Nevertheless, it is worth noting that there is still a 43.7% variance in the current dataset that cannot be explained by the linear model. Together, these data demonstrate that through-skull imaging can capture the mesoscopic features present in single-unit recordings, including the coarse tonotopic gradients, at least qualitatively.

Furthermore, to quantify the variability of tunings in the single-unit recording tracks, the standard deviation of all BFs along each track was calculated as a relative change to the mean BF of the track and shown on a logarithmic semitone scale (Figure 1E). The mean standard deviation across all tracks is 10.0 semitones across all subjects and is 10.0 (M60F), 9.2 (M160E), and 10.7 (M56E) semitones for each individual subject respectively (10 semitones ≈ 44% relative change). Although tracks close to the low-frequency reversal between A1 and R (hole 12 in M60F, hole 5 in M56E) seem to have a trend of being relatively more variable than the other tracks, the limited coverage of the current dataset hinders us from concluding on any specific spatial pattern of BF variability. Furthermore, the distribution of single unit tunings of each track in all three subjects is plotted in Figure 1F. A total of 164 tone-responsive tracks (rows) are sorted by their mean BFs (depicted by a black line). Each row shows a histogram of the single-unit tunings in that track. Notably, the spread of the single-unit tuning distributions remains relatively consistent across varying mean BFs.

### Comparison between tonotopic maps obtained by through-window intrinsic signal imaging under phase-encoded and trial-based experimental paradigms

In our previous work, the cortical responses imaged through-skull have been validated by demonstrating a high degree of similarity between the tonotopic map acquired through-skull and that acquired through a chronically implanted window (Song, et al., 2022). To further investigate the utility of intrinsic signal imaging for mapping tonotopy in awake marmosets with less attenuated signal, we measured tonotopy in a subject (M96B) through a round-shaped chronically implanted cranial window (Figure 2A) with two different experimental paradigms.

**Figure 2.**
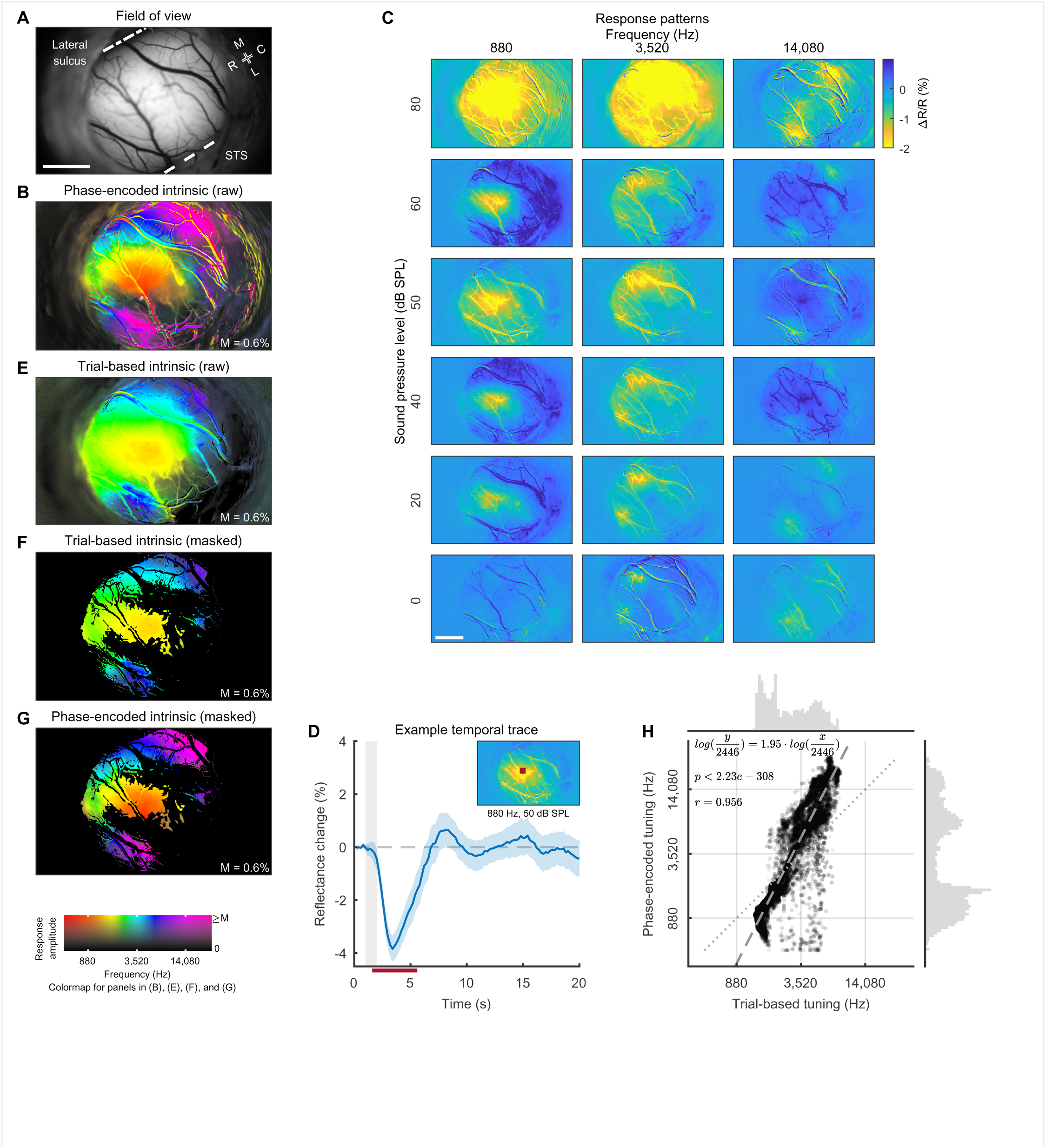
Comparison between tonotopic maps obtained with phase-encoded and trial-based paradigms. A. The field of view through the chronically implanted cranial window in the subject M96B. STS: superior temporal sulcus. Scalebar: 2mm. R: rostral; C: caudal; M: medial; L: lateral. B. The raw tonotopic map obtained with the phase-encoded paradigm. C. A trial-based experiment that measures the responses to pure tones of three different frequencies and at different levels. Response is quantified as the relative light intensity change averaged across the time window labeled by the red bar in (D). Scalebar: 2mm. D. The intrinsic optical signal trace of an exemplar pixel responding to a pure tone of 880 Hz delivered at 50 dB SPL. Blue solid line: mean response trace (n=20). Blue shade: standard error mean (n=20). Gray vertical shade: the stimulus duration. Red horizontal bar: time window over which the response pattern was averaged. Inset: the location of the exemplar pixel (denoted by a red dot). E. The raw tonotopic map estimated by the trial-based experiment in (C). F. The masked tonotopic map estimated from the trial-based experiment. The mask is a conservative estimation of pure-tone responsive pixels based on the following criterion: (1) both trial-based and phase-encoded response amplitudes are more than 0.3%; (2) p-value in the phase-encoded experiment <0.01. (3) the estimated hemodynamic delay is between 1.0 and 5.7 seconds. The same mask applies in (G). G. The masked tonotopic map estimated from the phase-encoded experimental paradigm. Colormap: the 2D color code for visualizing the tone-tuning frequency (color) and the response amplitude (intensity) together for each pixel. The upper display limit for response amplitude is labeled by “M”. The same colormap applies in (B), (E) and (F). H. Comparing the frequency tuning estimated from phase-encoded and trial-based experiments for the masked pixels (n = 24165). The masked pixels are shown as colored pixels in (F) and (G). Dotted line: a unity line. Dashed line: the linear fit of the tuning frequencies with a slope of 1.95 and an interception with the unity line at 2446 Hz (p-values < 2.23e-308). The correlation coefficient is 0.956. The histogram along the horizontal (vertical) axis shows the distribution of pixel tunings in the trial-based (phase-encoded) experiment.

The aforementioned tonotopy mapping was performed with a phase-encoded experimental paradigm, in which a tone pip sequence is played with increasing or decreasing frequencies continuously and repeated for twenty cycles. The response to the sound sequence is then Fourier-transformed to extract the frequency component corresponding to the sequence repetition. From the phase of this frequency component, the preferred frequency of each pixel can be derived (Song, et al., 2022). In the window-implanted subject (M96B), a tonotopy map estimated by this phase-encoded paradigm is shown in Figure 2B. Tonotopically organized regions corresponding to the auditory fields A1, R, and the recently described RPB high-frequency end are evident in the map. These results align with previous research that has demonstrated this phase-encoded paradigm as a powerful tool for mapping topographical organizations in the cortex with a substantial reduction in acquisition time and less susceptibility to artifacts such as heartbeats and respiratory motions (Kalatsky & Stryker, 2003) (Nelken, et al., 2004) (Kalatsky, et al., 2005) (Joly, et al., 2014) (Baumann, et al., 2015) (Song, et al., 2022). Nonetheless, it should be noted that the phase-encoded experiment we performed here cannot reveal functional details such as the response pattern to a particular frequency at a given sound level. A conventional trial-based experiment, on the other hand, measures neural response trial-by-trial and would thus provide complementary information and further validations to our phase-encoded experiment.

The neural responses to three pure tones at frequencies of 880, 3520, and 14080 Hz were measured at several different sound levels ranging from 0 to 80 dB SPL (Figure 2C). At each sound level, the three frequencies were delivered in a pseudo-randomized order within each of the 20 cycles. The intrinsic optical signal response to a frequency was then averaged across a response time window within the trial and across repetition cycles (Figure 2D). The response patterns to each frequency at levels above the hearing threshold (Osmanski & Wang, 2011) can be measured with clear responses (Figure 2C). The spread of the pattern generally increases with the sound level. When the sound level is high (e.g., 80 dB SPL), the spreads of the pattern at 880 and 3520 Hz expand almost across the entire optical window, whereas when the level is moderate (e.g., ∼50 and 60 dB SPL), the response pattern to each of the three tested frequencies is generally constrained within one or two specific parts of the optical window.

To estimate a trial-based tonotopic map, a “best frequency” for each pixel was computed by taking a weighted average of its responses to the three frequencies. The resulting tonotopic map is shown in Figure 2E. Although the same tonotopic gradients that appear in the phase-encoded map are also largely evident in the trial-based map, the estimated frequency range is more limited in the trial-based map. This can be at least partially explained by the stimulus difference between the two experiments: the phase-encoded stimuli cover 6 octaves, whereas the trial-based stimuli cover only 4 octaves. To further quantify the relationship between the two maps, pixels were selected by a mask that was based on response amplitude, statistical significance, and hemodynamic delay (Figure 2F, G and methods). The tunings of the selected pixels between the two paradigms were then compared in Figure 2H. The two measurements are highly correlated (r=0.956, p-value <2.23e-308, with a slope of 1.95), indicating that both methods can capture the major gradients of the tonotopic organizations but with some quantitative differences in estimated tuning. Nevertheless, the trail-based measure employed here has a longer acquisition time and still cannot cover the frequency as adequately as the phase-encoded measure.

### Comparison between tonotopic maps obtained by calcium imaging and intrinsic signal imaging

To further validate the tonotopic map using an alternative optical imaging modality, wide-field calcium imaging was performed in a GCaMP6s (Chen, et al., 2013) labeled subject (M80Z). The tonotopy was mapped with a trial-based experimental paradigm (Figure 3). Stimulus frequencies were sampled at each octave between 110Hz and 14080 Hz in this calcium imaging experiment (Figure 3A). This is much denser compared to the trial-based intrinsic signal imaging experiment presented in Figure 2C, owing to the much faster and more sensitive calcium signal (Figure 3B). The calcium signal was labeled by a dual-virus strategy with a combination of AAV1-CaMKII-cre and AAV(DJ)-FLEX-GCaMP6s (Song, et al., 2022). This strategy presumably labels calcium signals in excitatory neurons (Dittgen, et al., 2004), and thus measures neuronal activities more directly than the intrinsic optical signal. The intrinsic optical signal, on the other hand, reflects the hemodynamic changes following the local neural activities and is therefore slower and less sensitive than the calcium signal. The calcium response patterns at each frequency were evident across all levels above the hearing threshold (Osmanski & Wang, 2011). These patterns (Figure 3A) were also less susceptible to blood-vessel-related artifacts compared to the intrinsic signal patterns (Figure 2C). In terms of temporal dynamics, the calcium signal exhibits a rapid (∼0.3s) and positive onset comparing to the slow (∼2.4s) and negative intrinsic signal onset (Figure 3B). The faster speed of the calcium signal allows the trial length to be shortened by roughly four folds compared to the length of the intrinsic trials (Figure 2D) and therefore expedited the data acquisition process.

**Figure 3.**
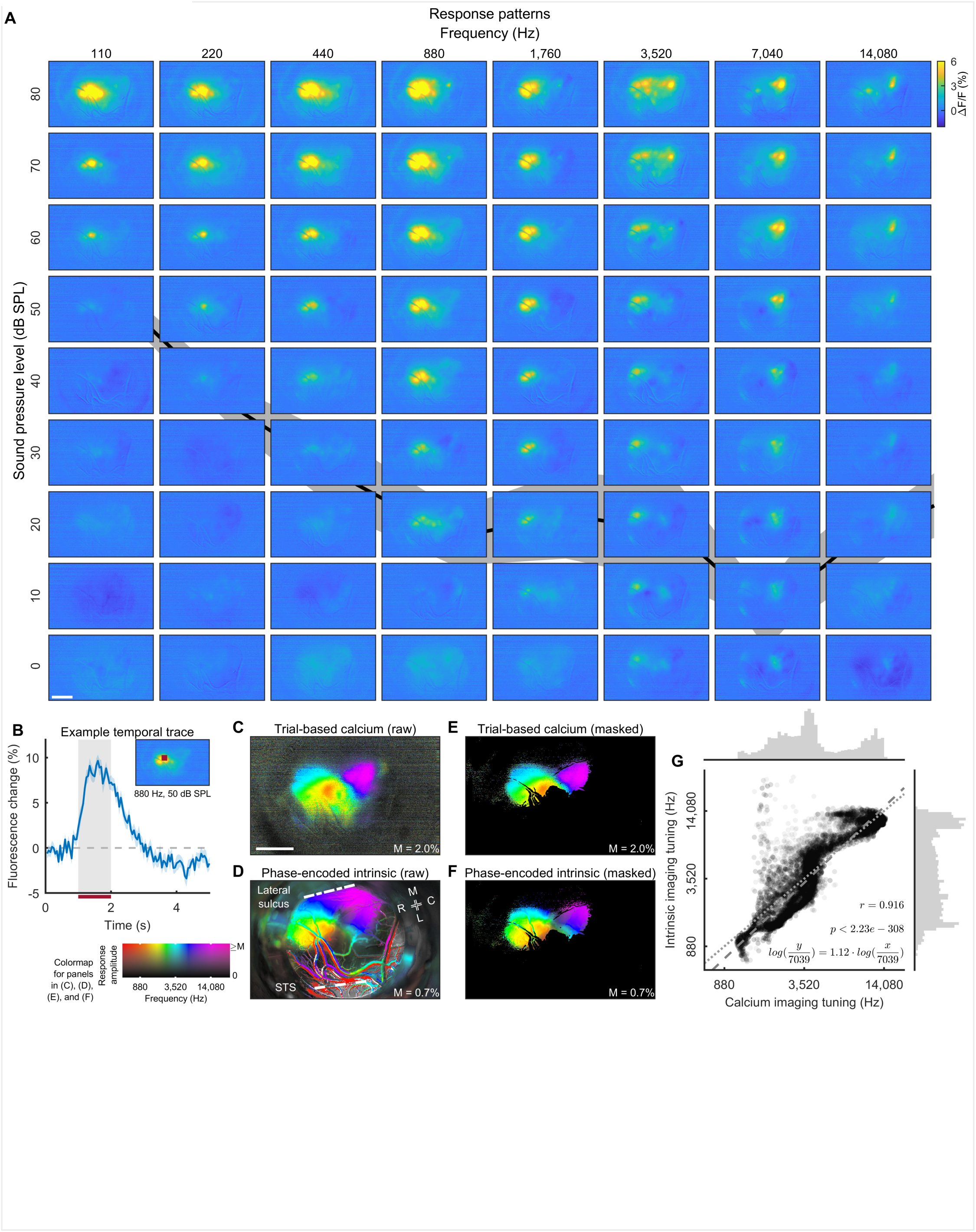
Comparison between tonotopic maps obtained by trial-based calcium imaging and phase-encoded intrinsic optical signal imaging. A. Calcium responses to pure tones of eight different frequencies and at different levels in the subject M80Z. The response is quantified as the relative fluorescence change to the pre-stimulus baseline. Scalebar: 2mm. The background curve and shadow show the mean and standard error mean of the marmoset hearing thresholds (Osmanski & Wang, 2011). B. The calcium signal trace of an exemplar pixel responding to a pure tone of 880Hz delivered at 50 dB SPL. Blue solid line: mean response trace (n=20). Blue shade: standard error mean (n=20). Gray vertical shade: the stimulus duration. Red horizontal bar: time period over which the response was averaged and shown in (A). Inset: the location of the exemplar pixel (denoted by a red dot). C. The raw tonotopic map estimated from the trial-based calcium imaging experiment. Scalebar: 2mm. D. The raw tonotopic map estimated from phased-encoded intrinsic signal imaging experiment. R: rostral; C: caudal; M: medial; L: lateral. STS: superior temporal sulcus. Colormap: the 2D color code for visualizing the tone-tuning frequency (color) and the response amplitude (intensity) together for each pixel. The upper display limit for response amplitude is labeled by “M”. The same colormap applies in (C), (E), and (F). E. The masked tonotopic map estimated from the trial-based calcium imaging experiment. The mask is a conservative estimation of pure-tone responsive pixels based on the following criterion: (1) Calcium response amplitude > 1%; (2) Phase-encoded intrinsic signal response amplitude > 0.1%; (3) p-value in the phase-encoded experiments <0.01; (4) Hemodynamic delay estimated by the intrinsic signal is between 1.0 and 5.7 seconds. The same mask applies in (F). F. The masked tonotopic map estimated from the phase-encoded intrinsic signal imaging. G. Comparing the frequency tuning of the masked pixels estimated from calcium imaging and intrinsic imaging (n = 16354). The masked pixels are shown as colored pixels in (E) and (F). Dotted line: a unity line. Dashed line: the linear fit of the tuning frequencies with a slope of 1.12 and an interception with the unity line at 7039 Hz (p-values < 2.23e-308). The correlation coefficient is 0.916. The histogram along the horizontal (vertical) axis shows the distribution of pixel tunings in the calcium (intrinsic) imaging experiment.

Following a similar procedure as in the last experiment (Figure 2), we estimated the best frequency for each pixel by computing the weighted average of frequency responses, resulting in a trial-based calcium tonotopy (Figure 3C). Due to the limited coverage of the imaging window, only the gradients corresponding fields A1 and a portion of R were captured, irrespective of the usage of the calcium signal (Figure 3C) or the intrinsic optical signal (Figure 3D). An image mask based on response amplitude, statistical significance, and hemodynamic delay was subsequently applied to both the trial-based calcium map (Figure 3E) and the phase-encoded intrinsic map (Figure 3F). Comparison of the two masked maps revealed a high correlation between the frequency tunings derived from the two imaging modalities (Figure 3G), with a correlation coefficient of 0.916 (p-value < 2.23e-308). A linear model fit between these two measurements yielded a slope of 1.12, indicating that the tonotopic maps acquired via both the calcium signal and the intrinsic optical signal were highly similar. It is worth noting that although 84.0% of the total variance can be explained by this simple linear model, there are certain discrepancies that cannot be explained by the current analysis and model (e.g., the non-linear relationship between the two at high-frequencies). Nevertheless, these results demonstrated a general consistency between the two imaging modalities and offered an additional verification of reliable measurements of the tonotopy in the marmoset auditory cortex.

### Tonotopic mapping through chronically implanted windows in additional subjects

Having validated our intrinsic optical signal imaging results with other recording modalities and experimental paradigms, we seek to map tonotopic organization in additional marmoset subjects (Supplementary Figure 1 and Figure 4). In our initial subjects (M80Z, presented in Figure 3; M96B, presented in Figure 2, and M132D), a 1/4-inch diameter optical window was implanted that covered a large proportion of the auditory cortex. These subjects (rows #1 to #3 in Figure 4) are ranked by how much the field of view covers the rostral part of the auditory cortex, as indicated by the tonotopy mapped through the window (Figure 4A-C). The response amplitude and frequency tuning are separately shown in Figure 4A and B, while a combined map is further shown in Figure 4C. Putative parcellations based on the tonotopy are exhibited in Figure 4D (see also methods).

**Figure 4.**
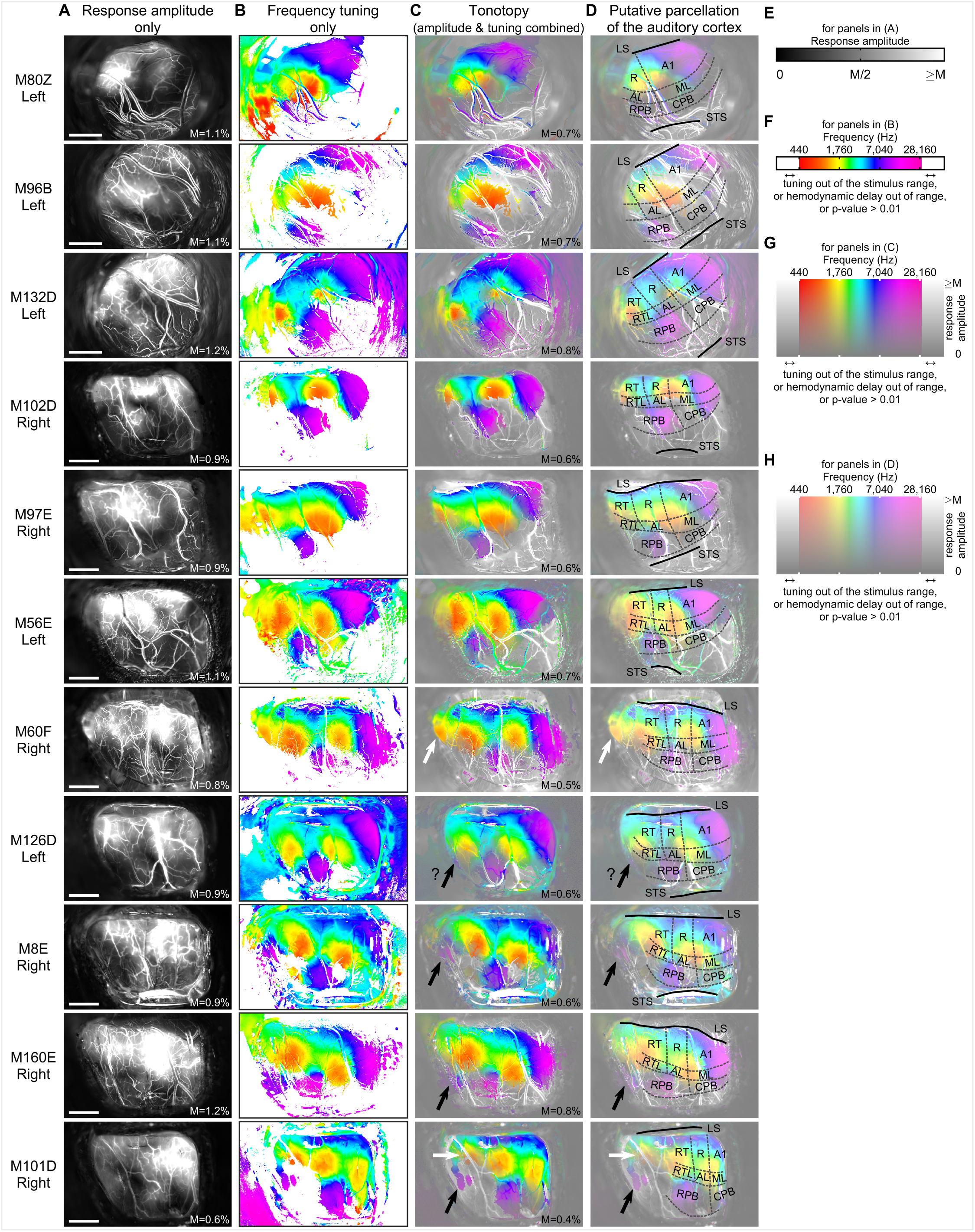
Tonotopic mapping through chronically implanted windows in more subjects. A. The response amplitude only map of each imaged hemisphere through a chronically implanted cranial window in awake marmosets. The subject ID and the imaged side are listed to the left of each panel. The right hemispheres are mirrored to the left for display purposes. The same mirroring fashion applies to all panels in this figure. “M”: the upper display limit of response amplitude (see also E). Scalebar: 2mm. B. The frequency tuning only map of each hemisphere. The tuning frequency is visualized as color for each pixel in these maps, according to the colorbar in (F). C. The combined tonotopy. The tuning frequency and response amplitude are visualized separately as color and intensity for each pixel in these maps, according to the 2D colormap in (G). The white (black) arrows indicate the locations of the low-(high-) frequency ends of a putative tonotopic gradient that is more rostral to RT, presumably in the rostrotemporal-polar field (RTp). The question marks indicate regions with ambiguity. The same labels apply in (D). “M”, the upper display limit of response amplitude, also applies in (D). D. The putative parcellation of the marmoset auditory cortex according to the tonotopic mapping results in (C). LS: lateral sulcus, STS: superior temporal sulcus, A1: primary auditory cortex (core), R: rostral core, RT: rostrotemporal core, ML: mediolateral belt, AL: anterolateral belt, RTL: rostrotemporal-lateral belt, RPB: rostral parabelt, CPB: caudal parabelt. E. The colorbar for panels in (A) F. The colorbar for panels in (B). For pixels with tunings out of the stimulus range, or hemodynamic delays out of range, or p-values > 0.01, the color saturation is reset to zero (white color). G. The 2D colormap for panels in (C). Estimated tone-turning frequency and response amplitude are coded in color (horizontal axis) and intensity (vertical axis), respectively. When an estimated tone-tuning frequency is not available (tuning phase out of the sound stimulus range), or the estimated hemodynamic delay is out of range, or with p-value > 0.01, the color is reset to grayscale. H. The 2D colormap for panels in (D).

Since this round window did not provide sufficient coverage of the entire auditory cortex on the marmoset brain surface, a larger trapezoid window was custom-made and implanted in subsequent subjects (Figure 4, row #4 and beyond). This window generally enabled imaging coverage over the entire extent of the auditory core (i.e., fields A1, R, and RT), the lateral belt (i.e., fields ML, AL, and RTL), and the parabelt (i.e., CPB and RPB). Not only the tonotopic gradients in the core, but also the recently described RT-RPB gradient (Song, et al., 2022) were visible through this window in all these subjects. More interestingly, in several subjects with the cranial window implanted even more rostrally (Figure 4, row #7 and beyond, subjects M60F, M126D, M8E, M160E, and M101D), additional tonotopically tuned regions that are located more rostrally to RT were observed. This phenomenon was particularly noticeable in subject M101D (Figure 4, the last row), in which a continuous low to high tonotopic gradient was observed at the location more rostral to RT. This gradient ran in the direction roughly perpendicular to the lateral sulcus, with its high-frequency end pointing more laterally. These observations prompted us to search more rostrally for unconventional tonotopically-organized-regions.

### Tonotopic mapping through un-thinned intact skull in additional subjects

Encouraged by the results in Figure 4, we extended the through-skull imaging FOV more rostrally in additional subjects (Supplementary Figure 2). The imaged tonotopic maps (Supplementary Figure 2 and Figure 5A-C) had better coverage of tonotopically organized regions on the rostral side than the through-window maps (Figure 4). A zoomed-in view showed this gradient rostral to RT can be observed in most of the hemispheres (Figure 5D) and confirmed the direction of this gradient runs roughly perpendicular to the lateral sulcus with its high-frequency end pointing more laterally. In contrast, the high-frequency end of the A1 gradient points more caudally whereas the high-frequency end of the RT-RPB gradient points roughly caudolaterally. The auditory areas can be putatively parcellated based on the tonotopic signatures as exhibited in (Figure 5E). Notably, the location of this additional rostral gradient (Figure 5F) was generally consistent with the recently identified primate auditory area “rostrotemporal polar” or “RTp” by cytoarchitecture and connectivity (Saleem, et al., 2007) (Saleem & Logothetis, 2012) (Scott, et al., 2017). This RTp region is at the rostral end of a caudorostrally directed serial cascade of connected auditory areas consisting of A1, R, RT, and RTp (Scott, et al., 2017).

**Figure 5.**
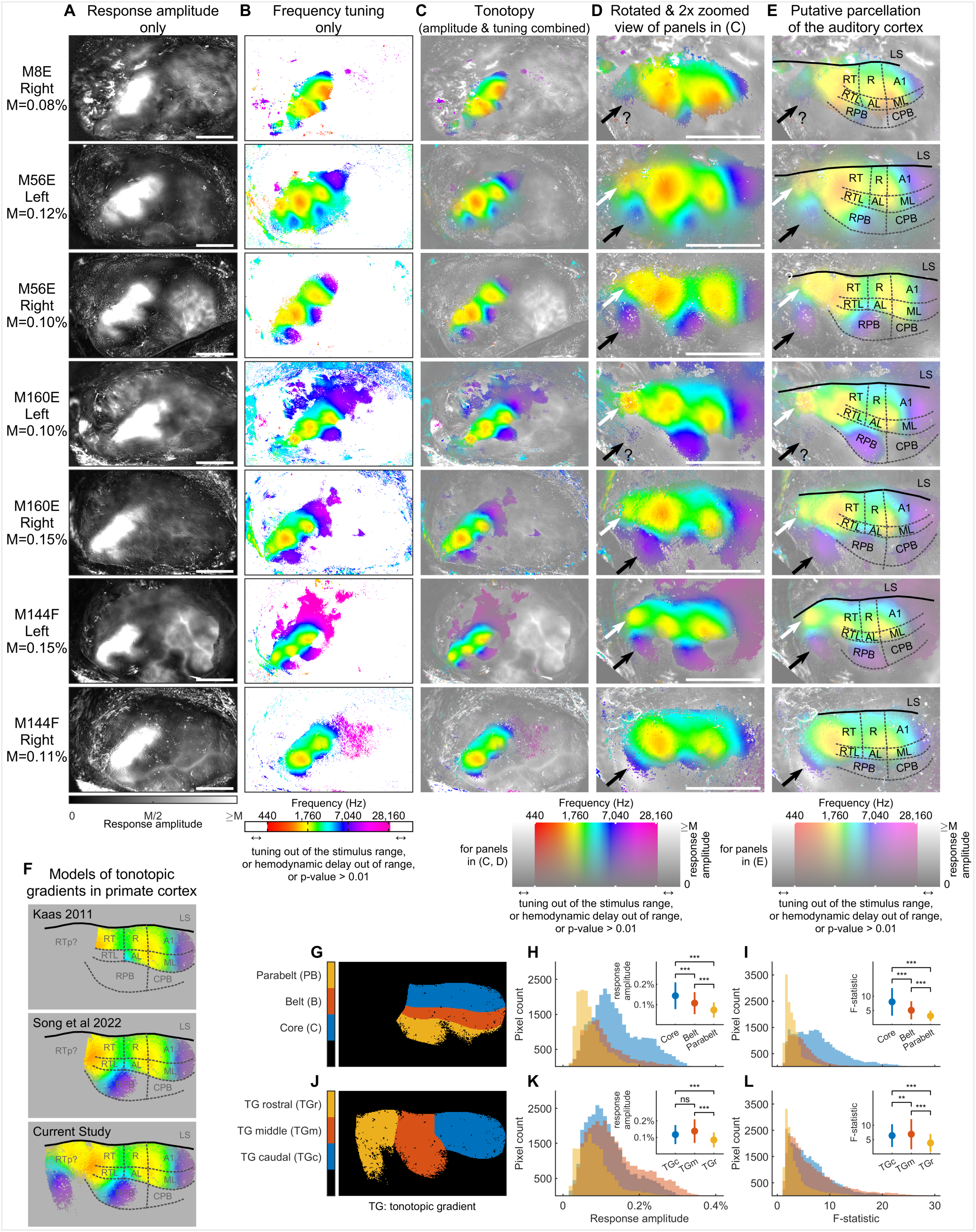
Tonotopic mapping through the intact un-thinned skull in more subjects. A. The response amplitude only map) of each imaged hemisphere through the intact un-thinned skull in awake marmosets. The subject ID, the imaged side, and the upper display limit of response amplitude “M” are listed to the left of each panel. The right hemispheres are mirrored to the left for display purposes. The same mirroring fashion applies to all panels in this figure. Scalebar: 5mm. B. The frequency tuning only map of each hemisphere. The tuning frequency is visualized as color for each pixel in these maps, according to the colorbar at the bottom. For a pixel, when an estimated tone-tuning frequency is not available (tuning phase out of the sound stimulus range), or the estimated hemodynamic delay is out of range, or p-value > 0.01, its color saturation is reset to zero (white color). C. The combined tonotopy. The tuning frequency and response amplitude are visualized separately as color and intensity for each pixel in these maps, according to the 2D colormap below. D. The rotated and 2× zoomed-in view of maps in (C). The white (black) arrows indicate the locations of the low-(high-) frequency ends of a putative tonotopic gradient that is more rostral to RT, presumably in the rostrotemporal-polar field (RTp). The question marks indicate regions with ambiguity. The same labels apply in (E). Scalebar: 5mm. E. The putative parcellation of the marmoset auditory cortex according to the mapping results in (D). LS: lateral sulcus, A1: primary auditory cortex (core), R: rostral core, RT: rostrotemporal core, ML: mediolateral belt, AL: anterolateral belt, RTL: rostrotemporal-lateral belt, RPB: rostral parabelt, CPB: caudal parabelt. F. The models of tonotopic gradients in primate cortex, including the models in (Kaas, 2011), (Song, et al., 2022), and current study. G. The parcellation of the auditory core (C), belt (B) and parabelt (PB) along the mediolateral hierarchy, using M56E’s right hemisphere as a schematic example. H. The response amplitude histograms of pixels in the auditory core (n=29383), belt (n=13227) and parabelt (15036) across all imaged hemispheres. The inset shows the mean ± standard deviation in each ROI. The same inset fashion applies in (I, K, and L). Along the hierarchy, the response amplitude is higher in any earlier tier than in a later tier (C vs B, B vs PB, C vs PB, p-values < 2.23e-308, Wilcoxon rank sum test, one-sided). I. The F-statistic histograms of pixels in the auditory core, belt and parabelt across all imaged hemispheres. Along the hierarchy, the F-statistic is higher in any earlier tier than in a later tier (C vs B, B vs PB, C vs PB, p-values < 2.23e-308, Wilcoxon rank sum test, one-sided). J. The parcellated ROIs of the tonotopic gradients: caudal (TGc), middle (TGm), rostral (TGr) along the mediolateral stages, using M56E’s right hemisphere as a schematic example. K. The response amplitude histograms of pixels in TGc (n=30578), TGm (n=31935) and TGr (n=16016) across all imaged hemispheres. Among these stages, the response amplitude is higher in TGc and TGm than in the newly identified TGr (TGc vs TGr, TGm vs TGr, p-values < 2.23e-308, Wilcoxon rank sum test, one-sided). L. The F-statistic histograms of pixels in TGc, TGm, and TGr across all imaged hemispheres. Among these stages, the F-statistic is higher in TGc and TGm than in the newly described TGr (TGc vs TGr, TGm vs TGr, p-values < 2.23e-308, Wilcoxon rank sum test, one-sided).

To further seek for functional differences among auditory regions, the response property is quantified and compared along both mediolateral and caudorostral stages. Figure 5G illustrates the region-of-interests (ROIs) parcellated along the mediolaterally directed auditory hierarchy (core, belt, and parabelt). The response amplitude is higher in the earlier stages than the later stages (Figure 5H), whereas the F-statistic (the ratio of variance explained by the stimulus over unexplained variance) is also higher in the earlier stages than the later stages (Figure 5I). These differences are consistent with the classic model of the primate auditory cortex (Kaas & Hackett, 2000) that narrowband sounds are better represented in the auditory core than in the auditory belt (Rauschecker, et al., 1995) (Rauschecker & Tian, 2004). The response property is also compared along the caudorostral direction. The extent of responsiveness along this direction can be divided into three major tonotopic gradients (TG): the caudal (TGc), the middle (TGm), and the rostral (TGr), each contains a complete and continuous representation of the entire frequency range (Figure 5J). Both TGc and TGm have higher response amplitude and F-statistic than TGr (Figure 5K, L) supporting the existence of a mediorostrally directed cascade for auditory information flow. These quantitative differences observed in the grouped results (Figure 5G-L) also hold in each and every individual hemisphere mapped (Supplementary Figure 3). With the verification analyses by other recording modalities and limitations discussed in the previous sessions (Figures 1, 2, and 3), these through-skull and through-window tonotopic maps (Figures 4 and 5) strongly support the existence of a tonotopic gradient rostral to RT that runs roughly perpendicular to the lateral sulcus in a medial-to-lateral direction, exceeding beyond the conventional auditory regions. Together, the newly identified rostral tonotopic gradient, joined with the recently described RT-RPB gradient (Song, et al., 2022) and the A1 gradient, form a periodic pattern along the caudorostral axis with at least 3 repeats of tonotopic gradients and extend our view on how primate auditory regions are tonotopically organized (Figure 3F).

## DISCUSSION

In this study, we used wide-field intrinsic optical signal imaging and calcium imaging techniques to map tonotopic organization in awake marmosets. The intrinsic optical signal imaging results obtained through-skull were validated by the data from window-implanted subjects. The intrinsic optical signal imaging results were cross-validated by wide-field calcium imaging and also electrophysiological recordings. The phase-encoded intrinsic signal imaging paradigm was also cross-validated by a trial-based paradigm with both intrinsic signal and calcium signal. These results demonstrated that the wide-field imaging techniques in awake marmosets provide a powerful platform for functional investigations at mesoscopic scales. Furthermore, we identified an additional tonotopic gradient in marmoset auditory cortex.

### New tonotopic gradients in primate auditory cortex

In addition to a recently reported new tonotopic gradient from RT to parabelt (Song, et al., 2022), we discovered another previously unknown tonotopic gradient that is located more rostral to RT (Figure 4 and 5). This gradient runs roughly perpendicular to the lateral sulcus, with its high-frequency end pointing more laterally (Fig. 4 and 5). The existence of such a gradient would call for a revision on our current view of how the primate auditory cortex is functionally organized (Kaas & Hackett, 2000) (Kaas, 2011) (Song, et al., 2022), especially along the caudo-rostral axis, a pathway that is hierarchically interconnected among several stages, including the recently identified area RTp (Saleem, et al., 2007) (Saleem & Logothetis, 2012) (Scott, et al., 2017).

The unexpected discovery of the tonotopic gradient rostral to the RT prompts several questions that remain unanswered. To name a few, it is unknown that how this functionally measured gradient aligns with the parcellations estimated based on other non-functional measurement modalities, such as cytoarchitecture mapping and connectivity mapping. Furthermore, the existence of a similar gradient in other primate species, such as macaques or humans, in which the auditory cortex is hidden within the lateral sulcus and other sulci, remains unknown. If such a gradient is present in these species, one might expect a relatively different anatomical orientation from that observed in marmosets due to factors like gyrification differences. In fact, it is plausible that other primate species also possess a similar gradient, as previous recording in the macaque STP suggested the existence of an additional tonotopic reversal that is rostral to RT using micro electrocorticogram (Fukushima, et al., 2012).

The existence of multiple tonotopic gradients along the rostro-caudal axis of the primate auditory cortex also poses a compelling question regarding their functional difference. It is currently unclear whether each tonotopic gradient possesses distinct functional specializations. One plausible hypothesis is that there exists a functional hierarchy from the caudal to rostral sub-fields: more rostral areas process information that is increasingly behaviorally relevant. A recent macaque study showed that identity selectivity of con-specific vocalizations generally increases along this caudal-to-rostral axis (Kikuchi, et al., 2010). Similarly, along the rostro-caudal axis of the temporal lobe, the ventral stream of primate visual cortex also has a mesoscopic repetition of topographically organized functional regions including the face patches (Bao, et al., 2020). The number of repetition cycles of the visual gradients seems matching that of the auditory gradients. Is there any co-alignment between the two? Is this type of periodic pattern of multiple functional gradients along the caudorostral axis a general principle of how primate temporal lobe is organized? All these questions may require future effort to answer.

### Comparison between mapping methods

#### Electrophysiology

In the realm of mesoscopic mapping of cortical functions in primates, electrophysiology has been the gold standard (e.g., auditory cortex mapping (Merzenich & Brugge, 1973) (Imig, et al., 1977) (Morel & Kaas, 1992)). However, the traditional approach of acutely prepared subjects comes with limitations. Although it provides accurate spatial information about recording tracks in the brain, the use of anesthesia may not sufficiently activate higher-order functional regions. Additionally, subjects have to be terminated after the procedure. In contrast, our electrophysiology preparation in awake marmosets involves a chronically implanted head-cap and an acutely inserted electrode for each recording session (Gao & Wang, 2020). This preparation allows us to record from the same marmoset subject for extended periods, ranging from months to years, while maintaining a recording density comparable to the acute preparation. It is worth noting that microscopic shifts between the brain and cranium can lead to less precise alignment of recording tracks across different days (∼<0.5mm), although mesoscopic functional features (∼>0.5mm) remain largely unaffected (Figure 1).

#### Electrophysiology vs. optical imaging

While electrophysiology provides a direct and reliable measurement of neural activities, it falls short in terms of efficiently covering a large proportion of the cortical surface. Consequently, its utility for mapping functional topography across multiple cortical areas at a mesoscopic scale is limited. Due to this efficiency constraint, our current electrophysiology dataset has a clear bias towards high-frequency regions since these data were mainly collected for another experiment and consumed for more than one year to collect. Furthermore, our current electrophysiology setup has certain configuration constraints that prevent electrodes from being lowered perpendicular to the local cortical surface in the most rostral part of the auditory cortex. This necessitates future modifications to enhance the capability of our electrophysiology coverage.

Optical imaging, on the other hand, offers an alternative technique that allows for more continuous coverage along the cortical surface. With the additional efforts we made to enlarge the FOV in Figures 4 and 5, more anterior and rostral regions can be recorded alongside the classical auditory cortex, thus expanding our ability to map functions in these regions simultaneously. Nonetheless, wide-field optical imaging techniques have their limitations as they primarily record structures on the brain surface. Consequently, the medial portion of the auditory core and the medial belt, which are concealed within the lateral sulcus, cannot be effectively captured. To overcome this limitation and enable comprehensive recordings encompassing the lateral auditory cortex, as well as the medial auditory areas and more anterior regions, further technical advancements are required.

#### Intrinsic optical signal imaging: advancements in through-skull and through-window approaches

In the past, optical imaging in primates was believed to be restricted to window-implanted subjects (Grinvald, et al., 1991) (Roe, 2007). However, we have recently developed a novel strategy that enables imaging of intrinsic optical signals through the intact un-thinned skull in awake marmosets (Song, et al., 2022). This through-skull intrinsic imaging method significantly expands our field of view for mapping cortical functions in marmosets (comparing Figure 5 to Figure 4) and serves as a guide for subsequent window implantation when higher imaging resolution is required. The results presented in Figure 5 highlight the effectiveness of this through-skull imaging approach, as it allows for coverage of more anterior and rostral areas and reveals additional functional gradients.

#### Experimental paradigms in intrinsic optical signal imaging

In the experiments presented in Figure 2, where we compared phase-encoded and trail-based paradigms, it is worth noting that the trail-based design only included three stimulus frequencies. This limited frequency sampling in the trial-based paradigm likely accounts for the high slope observed in Figure 2H. To obtain more accurate tonotopic gradients with the trial-based paradigm, it might be necessary to increase the density of stimulus frequencies and expand their range, as demonstrated in the calcium imaging experiment in Figure 3. However, denser frequency sampling in an intrinsic imaging experiment significantly extends the data acquisition time, which may not be ideal for mapping tonotopy. Nevertheless, these findings underscore the efficiency of phase-encoded intrinsic signal imaging as a valuable tool for mapping topographical organizations in the primate cortex (Joly, et al., 2014) (Baumann, et al., 2015) (Song, et al., 2022).

#### Calcium imaging

The tonotopy revealed by wide-field calcium imaging concurs with the results obtained through intrinsic imaging (Figure 3), further validating the utility of our optical mapping techniques. While the intrinsic optical signal exhibits slow dynamics (Figure 2D), the calcium signal is relatively fast and sensitive (Figure 3B), enabling sampling of a greater number of conditions within a single recording session. Additionally, the viral labeling strategy employed in our study (Song, et al., 2022) presumably expresses GCaMP mainly in cortical pyramidal neurons (Dittgen, et al., 2004), allowing for a more direct measurement of neuronal excitation compared to the hemodynamics-based intrinsic optical signal. Nonetheless, the current focus of wide-field calcium imaging is on functional organizations at mesoscopic scales, and it is plausible that additional organizational features exist at finer scales. This issue might be carefully addressed through the utilization of silent two-photon microscopy (Song, et al., 2022) as a potential solution.

## CRediT authorship contribution statement

Xindong Song: Conceptualization; Data curation; Formal analysis; Investigation; Methodology; Software; Validation; Visualization; Roles/Writing - original draft; Writing - review & editing

Yueqi Guo: Data curation; Formal analysis; Investigation; Methodology; Software; Validation; Visualization; Roles/Writing - original draft; Writing - review & editing.

Jong Hoon Lee: Data curation; Formal analysis; Investigation; Methodology; Resources; Software; Validation; Visualization; Writing - review & editing.

Chenggang Chen: Data curation; Investigation; Methodology; Validation; Writing - review & editing.

Xiaoqin Wang: Conceptualization; Funding acquisition; Project administration; Resources; Supervision; Writing - review & editing.

## Acknowledgement

This research was supported by National Institutes of Health grants DC003180 to X.W. X.S. was supported by a fellowship from the Kavli Neuroscience Discovery Institute at JHU. We thank J. Lynch, K. Schonvisky, S. Miller, E. Easter, and J. Izzi for assistance with surgeries and animal care, A. W. Roe for sharing cranial window related protocols.

## METHODS

### Animal preparation

Twelve marmosets were used in the current study. All twelve subjects were used in the tonotopic mapping experiment with intrinsic signal and phase-encoded paradigm (four subjects and seven hemispheres for through-skull imaging, eleven subjects and eleven hemispheres for through-window imaging). One subject (M96B) was used in the tonotopic mapping experiment with intrinsic signal and trial-based paradigm. One subject (M80Z) was used in the tonotopic mapping experiment with calcium signal and trial-based paradigm. Three subjects (M60F, M160E, M56E) were recorded with electrophysiology after through-skull imaging. The basic design of the marmoset chronic head-cap implant (Gao & Wang, 2020), as well as the optical imaging setup (Song, et al., 2022) has been described previously. Briefly, the head-cap for head fixation was implanted during an aseptic surgery. The entire skull was covered with a thick layer of dental cement, except the lateral part of the skull over the auditory cortex, which was covered with a thin layer (∼1 mm) of dental cement and formed an imaging chamber. The subjects were adapted to sit calmly in a Plexiglass restraint chair in an upright position head-fixed. The auditory cortex is imaged either through the intact skull, or through an artificial dura-based imaging window with the XINTRINSIC system (Song, et al., 2022). The imaging objective pointed from the lateral side of the animal towards the imaging chamber, along the direction perpendicular to the skull surface over the putative auditory cortex. All experimental procedures conformed to local and US National Institutes of Health guidelines and were approved by the Johns Hopkins University Animal Use and Care Committee.

### Auditory stimulus delivery

All experiments were conducted in a double-wall acoustic chamber (IAC acoustics, customized, ∼3.4 m (W) x 3.1 m(D) x 2.6 m (H) in size). The interior was covered with 3’’ acoustic foam (Pinta acoustic, SONEX). For auditory stimulation, digital waveforms (16-bit resolution, 100,000 samples/second conversion rate) were converted into analog signals on a NI DAQ card (NI PCIe-6323) that also controlled the imaging acquisition. The analog signal was attenuated by a programmable attenuator (Tucker-Davis Technology, PA5) before feeding into an amplifier (Crown, D75) that powered a loudspeaker (KEF, LS50), placed ∼1 m away in front of the subject. Sound levels were calibrated by placing a microphone (Brüel & Kjær, type 4191) at the animal’s typical head position.

### Tonotopic mapping by intrinsic imaging with a phase-encoded experiment paradigm

To map the tonotopy with intrinsic signal, a pure tone pip sequence with either ascending or descending frequencies was played for 20 repetitions (Song, et al., 2022). The sequence consisted of 73 pure tone pips in the middle of a 20-second trial, each pip with a 0.2-second duration and 20-millisecond sine ramps at both onset and offset. All pure tone pips were delivered at 50 dB SPL level, measured at the position of the animal’s head. The “ascending” sequence started with a pure tone of 440 Hz, continuing with each of the following pips ascending one semitone in frequency from the previous pip, and ending with a pure tone pip of 28160 Hz. The “descending” sequence was in the reversed order of the “ascending” sequence. The imaged responses were then Fourier-transformed, and the phase of the 0.05 Hz frequency component (corresponding to the 20-second repetition cycle) was extracted to estimate the tuning for each pixel. A pure-tone responsive tonotopic map was therefore derived by this “phase-encoded” experiment paradigm, which was described in a previous work (Song, et al., 2022) in detail.

### Tonotopic mapping by intrinsic imaging with a trial-based experiment paradigm

Trial-based intrinsic imaging was performed in subject M96B. Each trial started with 1-second of pre-stimulus silent period, followed by a 1-second-long pure tone with 20-millisecond sine ramps at the onset and the offset, and ended with an 18-second post-stimulus silent period for the hemodynamic response to settle. Three frequencies, 880, 3520, and 14080 Hz were played at a pseudorandom order for 20 repetitions at sound levels of 0-, 20-, 40-, 50-, 60-, and 80-dB SPL, while intrinsic responses were imaged.

To analyze the data from a trial-based experiment, the imaged light “reflectance” during the pre-stimulus period was averaged as a baseline (R). The change in the light intensity across each single trial was divided by the baseline (ΔR/R). The time of the response peak was estimated to be around 3.6 seconds after the stimulus onset based on the mean temporal trace across all pixels and all trials. The time integration window was empirically determined to be 4 seconds around this peak (1.6 seconds to 5.6 seconds from the trial start). The average change (ΔR/R) within this window was used as a measurement of evoked response. The signal has a negative polarity that a negative change in intensity means more absorption of the light, thus an increase in hemodynamic inflow. The response level of each pixel is plotted in Figure 2C to show the distinct response patterns to tones played at different frequencies and levels.

The best frequency of each pixel was estimated following the steps described below. For each frequency, the response patterns to three sound levels (40-, 50-, and 60-dB SPL) were averaged first. Next, for each individual pixel, its response amplitudes to different frequencies were sorted in descending order by their signal-polarity compensated values. The third (last) value in the order was used as a reference value and was deducted from all values, resulting in three differential values with the third equal to 0. The next step involved normalizing these differential values so that the weights for each pixel summed up to one. Finally, we applied these calculated weight values to the corresponding stimulus frequencies on a logarithmic scale to derive the best frequency estimation for each pixel. The best frequency values were shown following the same fashion as in the phase-encoded experiment: the hue (H channel in the HSV color space) in the map represents the best frequency, while both the saturation and value (S and V channels in the HSV color space) represent the maximum response to the three pure tones.

### Tonotopic mapping by calcium imaging with a trial-based experiment paradigm

A combination of two viral vectors carrying genetically encoded calcium indicators (AAV(DJ)-EF1a-DIO-GCaMP6s + AAV1-CaMKII0.4-Cre-SV40) were stereotaxically microinjected into the cortex through the artificial dura by six tracks to cover the auditory cortex within the window (Song, et al., 2022). 30 days were allowed for GCaMP to be expressed before calcium imaging experiments were performed.

In these experiments, eight frequencies were chosen from 110 to 14080 Hz with a step of one octave and were played at nine sound levels from 0 to 80dB SPL with a 10dB SPL step, resulting in a total of 72 conditions. The frequencies were played in a pseudo-random order and 20 repetitions were recorded for each condition. Each trial started with a 1-second pre-stimulus silent period, followed by a 1-second pure tone with 20-millisecond sine ramps at the onset and the offset, and ended with a 3-second post-stimulus silent period.

To analyze the data from calcium imaging, the averaged fluorescence level during the pre-stimulus period were used as a baseline (F), and the mean relative fluorescence change (ΔF/F) during the sound-presenting window were used as a measurement of the evoked response. The response of each pixel is plotted in Figure 3A to show the distinct response patterns to tones played at different frequencies and levels. The tonotopic map was plotted in the same way as trial-based intrinsic imaging (for M96B), with the exception that the deducted reference value was estimated as the average value among the third to the last sorted response amplitudes in the descending order, and the resulting differential response values were rectified before converting them into weight values.

### Tonotopic mapping by electrophysiological recording

Electrophysiology experiments were conducted inside a double-walled soundproof chamber (Industrial Acoustics, custom model), whose interior was covered by 3-inch acoustic absorption foam (Pinta Acoustic, SONEX). Neural recordings from auditory cortex of awake marmosets were acquired using a 64-channel probe (Model P64-12, Diagnostic Biochips. Inc.). After through-skull imaging, neural recordings were carried out in awake condition by inserting the 64-channel probe perpendicular to the cortical surface through a miniature craniotomy hole (diameter ∼1mm) drilled through the skull using previously described procedures (Gao & Wang, 2020). Between 5 to 10 recording tracks were made in each hole. Signals were recorded from the 64 active channels of the probe spanning across 640um in depth. Data were acquired by OpenEphys data acquisition board and software at 30kHz, with an amplifier gain of 192 for each channel and band-pass filtered with cutoff frequencies at 300 and 6000Hz. Spike sorting was performed using an automated spike-sorting Matlab interface (Kilosort) to cluster units from raw data (Pachitariu, et al., 2016). The resulting spike clusters were manually curated to split and merge clusters using Phy Template-GUI software (Kwik team). Further quality control was performed using custom Matlab code to ensure single-unit data. In total, 1955 single units were recorded across 99 tracks in 16 holes in animal M60F, 1211 single units across 60 tracks in 7 holes in animal M160E, and 1077 single units across 40 tracks in 5 holes in animal M56E. For subjects M160E and M56E, only the high frequency -regions were targeted for electrophysiology recording guided by the tonotopic map from through-skull imaging.

To characterize best-frequency tunings of recorded single-units, during each recording track, pure tone testing trials were delivered in a randomized order to the subject at the sound level of ∼60 dB SPL, near to our through-skull imaging testing level (50 dB SPL). Each pure tone trial contained a 200-millisecond pre-stimulus duration, a 100-millisecond stimulus duration, and a 300-millisecond post-stimulus duration. Pure tones were synthesized at a density of 10-20 steps/octave spanning up to 5 octaves to cover the track’s receptive field.

Individual peri-stimulus time histograms (PSTH) in response to pure tones were obtained by convolving a Gaussian kernel (σ = 20 millisecond) with the spike train. The position of a peak in the PSTH was found during the stimulus and post-stimulus duration periods. Only neurons with peak firing rates of two standard deviations above the mean spontaneous firing rate and an average firing rate of more than 1 spike/second were considered as having a pure-tone response. The corresponding position of the peak was used to define the best-frequency of the unit.

### Comparison between tonotopic maps obtained by different recording methods

To compare the frequency tunings obtained by intrinsic imaging and electrophysiological recording, we first aligned the positions of the recording tracks and craniotomy holes with the tonotopic maps measured by intrinsic imaging, using the fine structures in the recording chamber as landmarks. For track-wise comparison, we averaged the best frequencies of all units within each track, then compared it with the tuning of the corresponding pixel from imaging (Figure 1C). For hole-wise comparison, we averaged the best frequencies of all units recorded in the craniotomy hole, then compared it with the mean optically measured tuning averaged across all pixels within the hole (Figure 1D). A linear model was fit to each condition and was plotted as a dashed line together with the scatter plot.

To quantify the similarity between two tonotopic maps obtained by phase-encoded paradigm and trial-based paradigm (Figure 2), or calcium imaging and intrinsic imaging (Figure 3), we first performed non-rigid registration with NoRMCorre algorithms to align the two field of views by their anatomical features (Pnevmatikakis & Giovannucci, 2017), and then performed a linear model fit for a subset of pixels. These pixels were selected according to several response properties. In the phase-encoded intrinsic imaging experiments, a qualified pixel must have (1) a response amplitude larger than 0.3% (Figure 2, M96B) or 0.1% (Figure 3, M80Z), (2) an estimated hemodynamic delay between 1 second to 5.7 seconds, (3) a response profile that was significantly modulated at the repetition frequency 0.05 Hz in the phase-encoded intrinsic imaging experiment (one-way ANOVA, p < 0.01). When compared with trial-based intrinsic tonotopic mapping, an additional criterion includes a trial-based intrinsic response magnitude larger than 0.3% (Figure 2, for M96B). When compared with calcium signal tonotopic mapping, an additional criterion includes a calcium response magnitude larger than 1% (Figure 3, for M80Z). The resulting pixel-wise frequency tuning comparison were plotted in Figure 2H and Figure 3G, and linear models were fit to the data points, plotted as dashed lines together with the scatter plots.

### Parcellation of the auditory areas

The auditory cortical fields are defined functionally in the current study to maximize the consistency with the previously established convention for parcellating the primate auditory cortex [Kaas 2011]. The criteria are summarized and listed as follows:

1. Along the medial-lateral axis, the primate auditory cortex can be parcellated into three tiers: the auditory core (primary), the auditory belt (secondary), and the auditory parabelt (tertiary) by a series of boundary lines running roughly along the rostral-caudal direction.

a. The boundary between the core and the belt at its rostral proportion is defined by a line connecting the centers of the two caudal-most low-frequency reversals.
b. The boundary between the belt and the parabelt at its caudal proportion is defined by a line that marks where the tonal responsiveness starts to disappear
c. The core-belt and belt-parabelt dividing boundaries are extended from the proportions defined in (a) and (b) and are roughly parallel to each other at each proportion.
d. The parabelt outline boundary at its rostral proportion is defined by a line that marks where the tonal responsiveness ends, with the lateral high-frequency responsive region within the parabelt.
e. The caudal proportion of this parabelt outline boundary is roughly symmetric to its rostral counterpart.
2. Along the caudal-rostral axis, the core, the belt, and the parabelt can be each parcellated into several subfields by a series of medial-laterally running boundaries.

a. The auditory core can be parcellated into three subfields by two boundaries. One at the caudal-most low-frequency reversal separates A1 and R (rostral field), whereas the other one at the slightly more rostral high-frequency reversal separates R and RT (rostro-temporal field).
b. These two boundaries can be extended laterally to further parcellate the lateral belt into subfields ML (middle-lateral field), AL (anterior-lateral field), and RTL (rostro-tempo-lateral field), with ML parallel to A1, AL parallel to R, and RTL parallel to RT.
c. The more caudal boundary can be further extended laterally to divide the parabelt into CPB (caudal parabelt) and RPB (rostral parabelt).

