## Supplementary figures and images for "Tonotopic organization of auditory cortex in awake marmosets revealed by multi-modal wide-field optical imaging"

### Supplementary figures

#### Supplementary Figure 1

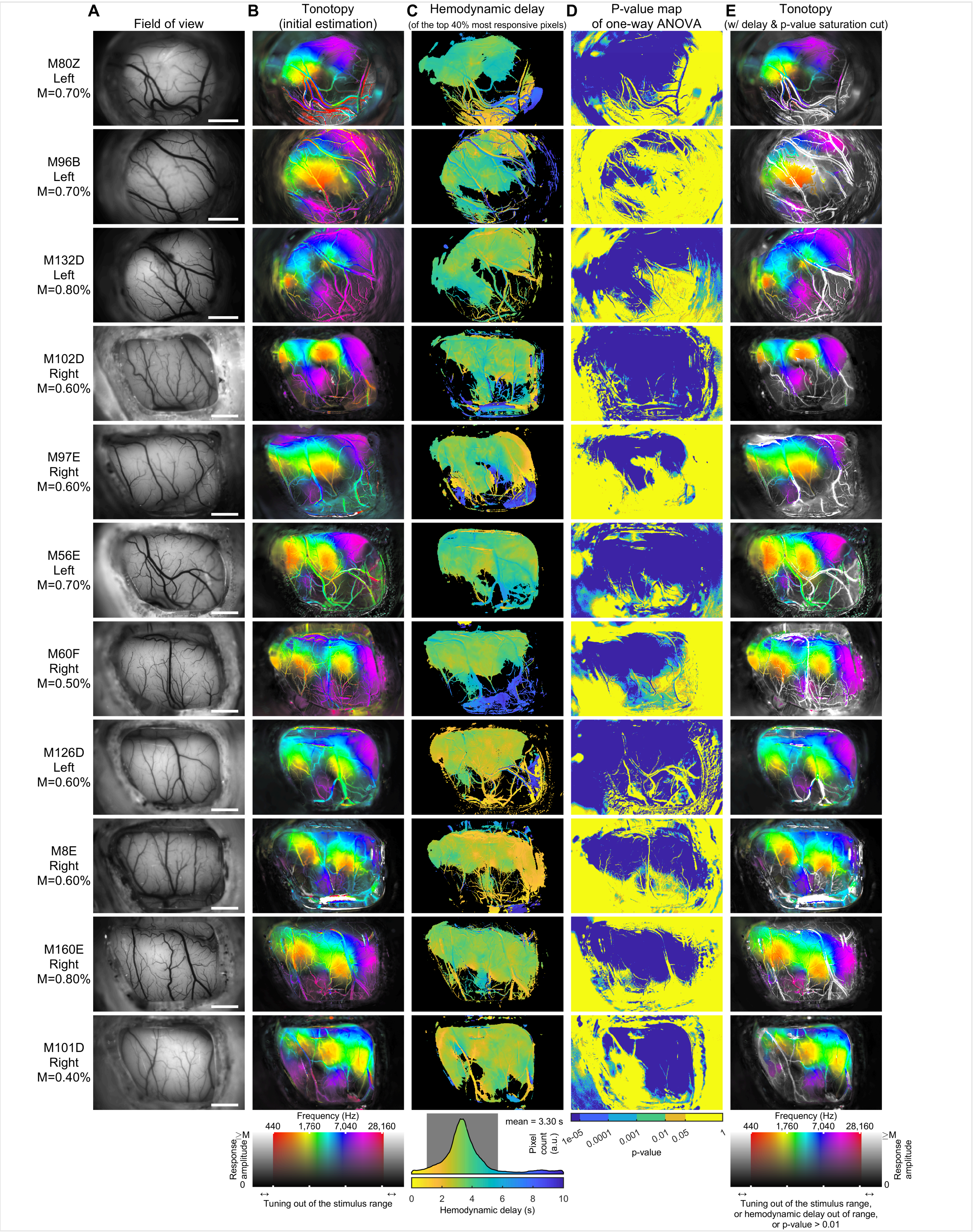

Supplementary Figure 2

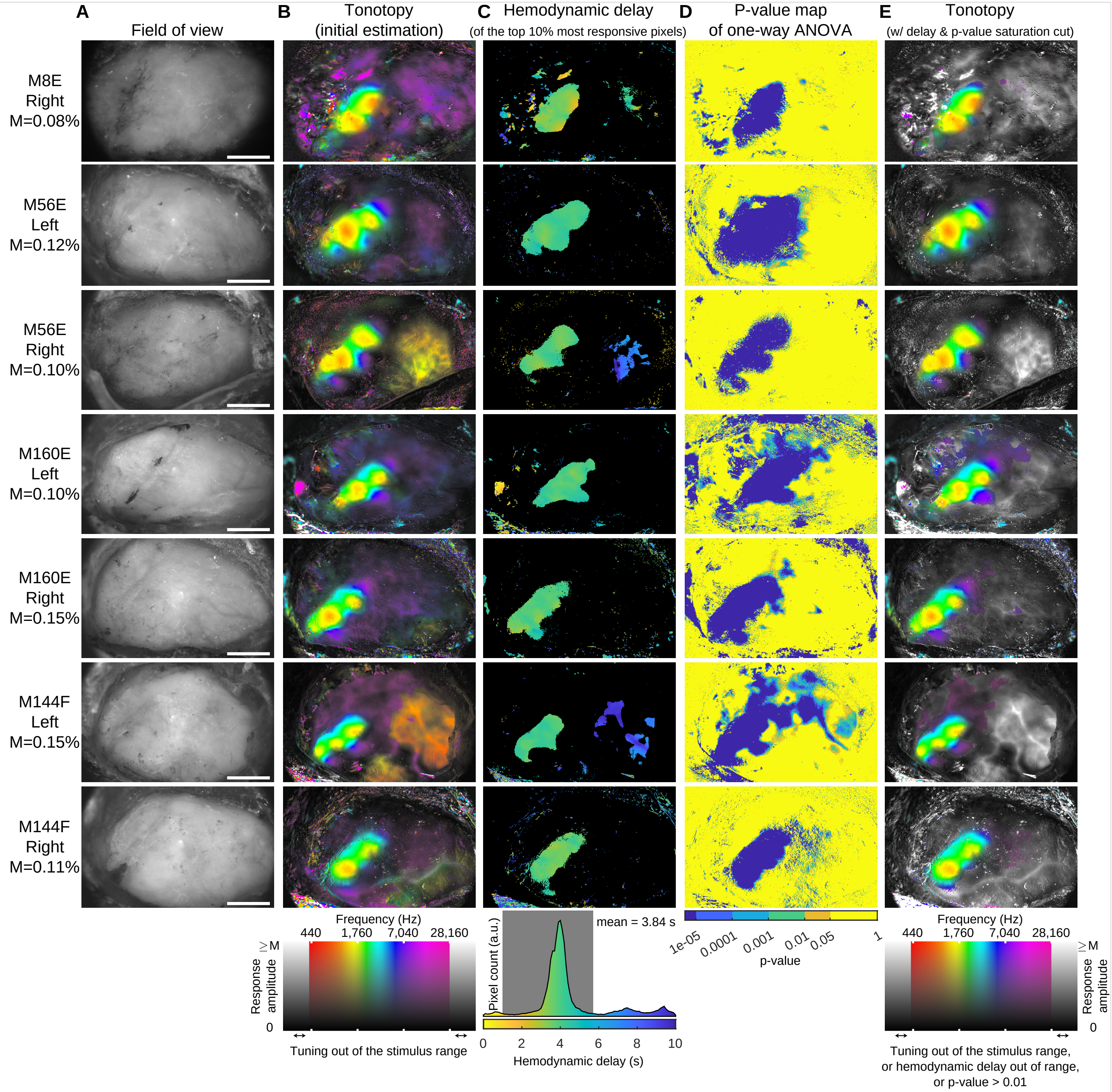

#### Supplementary Figure 3

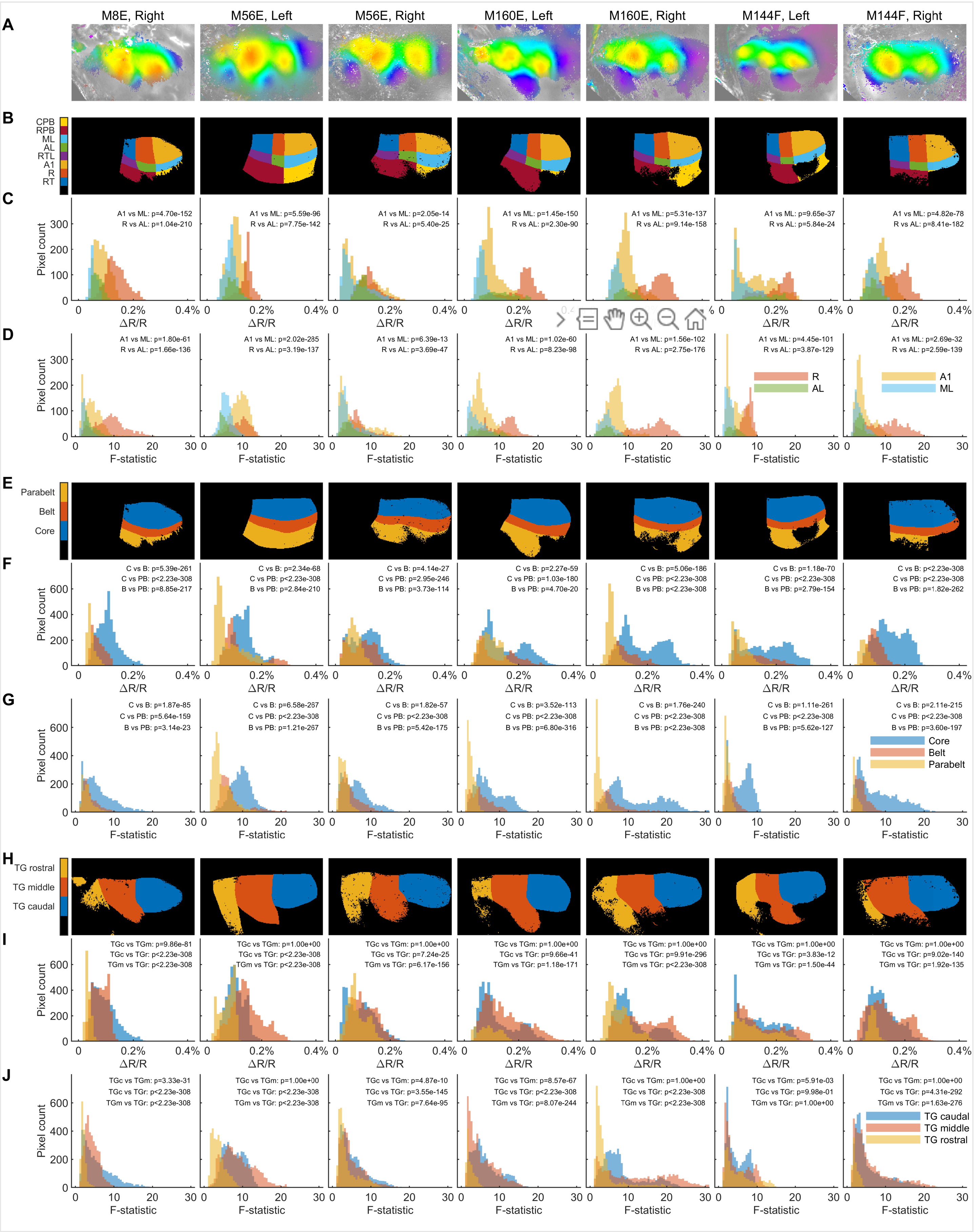
